# A curated transcriptome dataset collection to investigate inborn errors of immunity

**DOI:** 10.1101/526004

**Authors:** Salim Bougarn, Sabri Boughorbel, Damien Chaussabel, Nico Marr

## Abstract

Primary immunodeficiencies (PIDs) are a heterogeneous group of inherited disorders, frequently caused by loss-of-function and less commonly by gain-of-function mutations, which can result in susceptibility to a broad or a very narrow range of infections but also in inflammatory, allergic or malignant diseases. Owing to the wide range in clinical manifestations and variability in penetrance and expressivity, there is an urgent need to better understand the underlying molecular, cellular and immunological phenotypes in PID patients in order to improve clinical diagnosis and management. Here we have compiled a manually curated collection of public transcriptome datasets mainly obtained from human whole blood, peripheral blood mononuclear cells (PBMCs) or fibroblasts of patients with PIDs and of control subjects for subsequent meta-analysis, query and interpretation. A total of nineteen (19) datasets derived from studies of PID patients were identified and retrieved from the NCBI Gene Expression Omnibus (GEO) database and loaded in GXB, a custom web application designed for interactive query and visualization of integrated large-scale data. The dataset collection includes samples from well characterized PID patients that were stimulated *ex vivo* under a variety of conditions to assess the molecular consequences of the underlying, naturally occurring gene defects on a genome-wide scale. Multiple sample groupings and rank lists were generated to facilitate comparisons of the transcriptional responses between different PID patients and control subjects. The GXB tool enables browsing of a single transcript across studies, thereby providing new perspectives on the role of a given molecule across biological systems and PID patients. This dataset collection is available at: http://pid.gxbsidra.org/dm3/geneBrowser/list.

## Introduction

Primary immunodeficiencies (PIDs) are a heterogeneous group of inherited disorders, most often caused by loss-of-function mutations and less commonly by gain-of-function mutations, affecting components of the innate and/or adaptive immune system [1-3]. These inborn errors of immunity can result in profoundly increased susceptibility to a broad or a very narrow range of infections but also autoimmune disorders, allergies and malignancies [1, 4-6]. The spectrum of clinical manifestations of PIDs is very broad and largely dependent upon the affected gene(s) and the degree to which normal gene function is lost or altered. In addition, a variety of other factors such as germline or somatic mosaicism, modifier genes and environmental factors can play an important role in the clinical penetrance and expressivity of a given disease phenotype [3, 7, 8]. To date, mutations in more than 300 genes have been identified to cause PIDs, which are classified into major groups reflecting the diverse immunological phenotypes [5, 9]. Nonetheless, PIDs often go unrecognized or are not properly diagnosed [10]. However, with the recent developments and rapidly declining costs of next-generation sequencing technologies and other high-throughput methods, it is expected that many more as yet unknown disease-related genetic variants will be discovered in the near future [7].

A considerable challenge for identifying causal genetic variants—which is critical for the diagnosis and clinical management of PID patients—lies in the vast heterogeneity of the underlying immunological phenotypes and clinical manifestations on the one hand and in the degree of human genetic variation between individuals on the other hand. Despite considerable advances in recent years, specific gene functions in humans, their roles and regulation in biological processes, and essentiality of the redundancy in a function of a particular gene for protective immunity of the human host remains poorly understood [6]. In many cases, the use of forward and reverse genetics in mice or other model organisms has provided insufficient insights into the pathophysiology of PIDs due to interspecies differences and the fact that inbreeding has let to various deficiencies in laboratory animals, rendering them susceptible to a broad range of infections that often poorly recapitulates the clinical phenotypes in humans [11]. On the other hand, studying naturally occurring genetic defects in humans is much more challenging given the difficulty of obtaining biological samples, ethical implications and potential risks that go along with it. An additional challenge is the low frequency of most null alleles. Although PIDs are not necessarily rare when considered collectively, the small number of individuals that suffer from a specific deficiency usually does not permit classic case-control or family-based genetic association studies. Indeed, a considerable proportion of monogenic etiologies of PIDs were initially reported in single patients [12]. The ability to identify single-gene inborn errors in PID patients requires validation of the disease-causing variant by in-depth mechanistic studies demonstrating the structural and functional consequences of the mutations using blood or other accessible biological samples such as fibroblasts from skin biopsies [12]. In this context, several transcriptomics studies have been conducted using whole blood, PBMCs and fibroblasts of well-characterized PID patients, to assess the underlying immunological phenotypes at the molecular and cellular levels in more detail (table 1) and in many cases, to further validate the causal relationship between the underlying genotypes and clinical phenotypes. Notable are several seminal studies of PID patients with susceptibility to a very narrow range of pathogens, such as patients with MYD88 or IRAK4 deficiency who are primarily susceptible to pyrogenic bacterial infections [13, 14], patients with TBK1, TRIF or TLR3 deficiency [15-17] which underlies herpes simplex encephalitis of childhood, or a recent study of a child with IRF7 deficiency who was primarily susceptible to severe influenza but otherwise immunocompetent with regard to other common infectious diseases [18]. Such studies have highlighted that often, the underlying gene defect may only affect a narrow repertoire of transcriptional responses while the affected individual’s cells remain highly responsive to specific stimulation through alternate receptors, pathways and signaling networks and in particular to *ex vivo* stimulation with whole organisms, reflecting the high degree of human gene redundancy in host defenses [6].

**Table 1:**
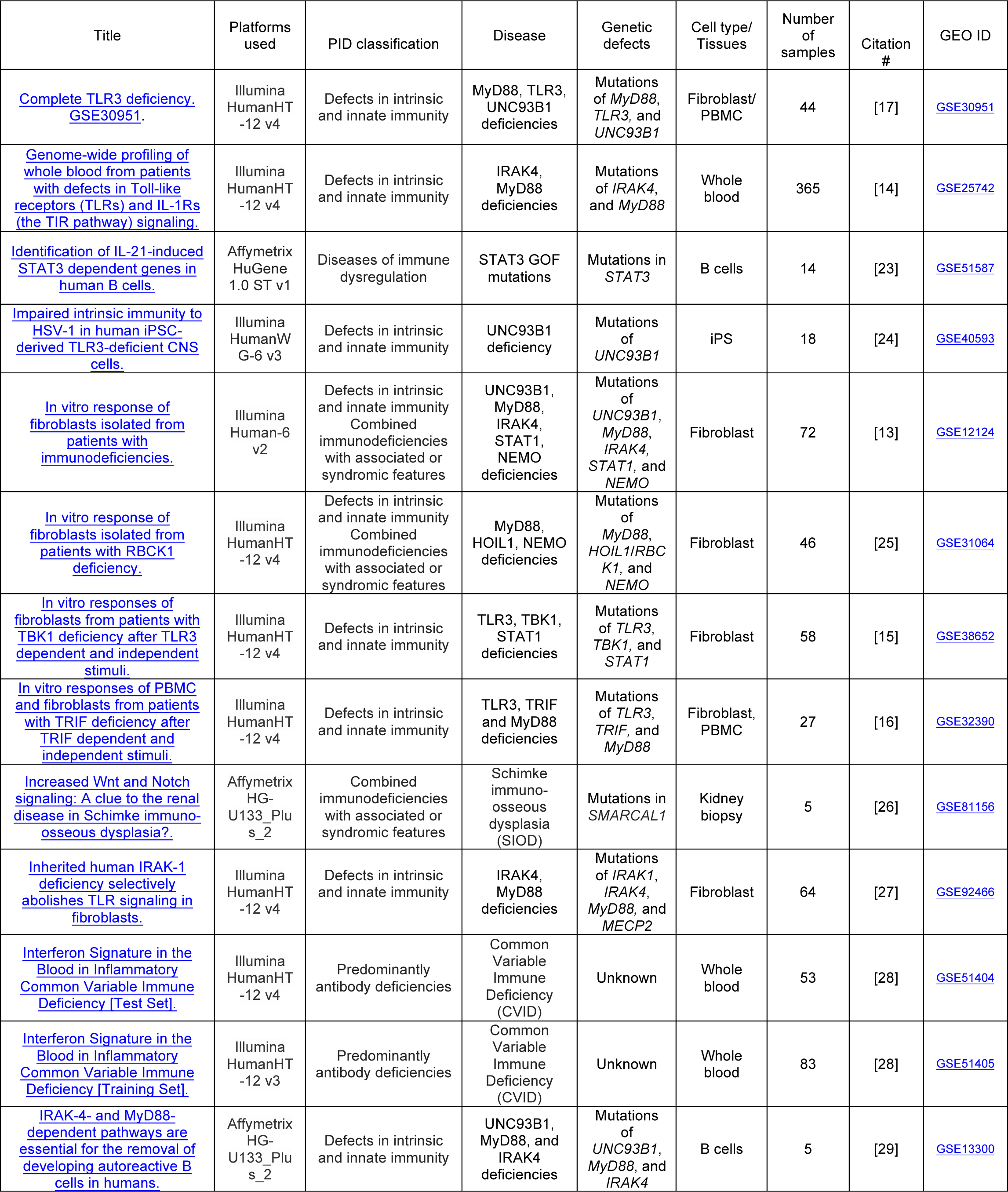
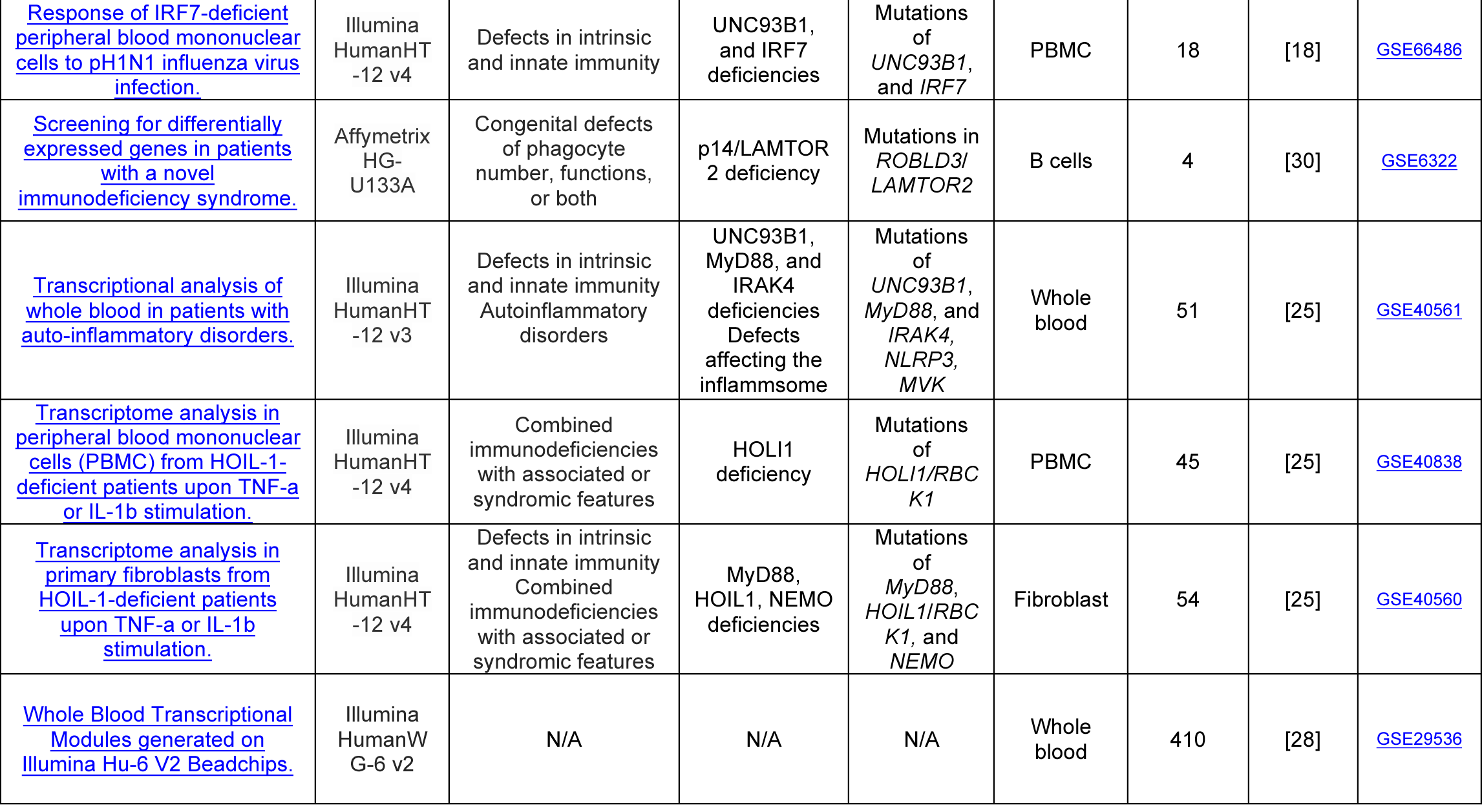
List of datasets constituting the collection.

Here, we compiled a curated collection of 19 transcriptome datasets, retrieved from the NCBI’s Gene Expression Omnibus (GEO) database, to provide as resource for the investigation on inborn errors of immunity. The datasets were loaded into a custom interactive web application, the Gene Expression Browser (GXB), (http://pid.gxbsidra.org/dm3/geneBrowser/list), which allows seamless access to the data and interactive visualization of the transcriptional responses, along with demographic and clinical information [19]. The user can customize data plots by adding multiple layers of parameters (e.g. age, gender, sample type and type of genetic defects), select and modify the sample ordering and gene rank lists, and generate links (mini URL) that can be shared via e-mail or used in publications. The GXB tool enables browsing of a single transcript across multiple studies and datasets, providing new perspectives on the role of a given molecule across biological systems and PID patients. In summary, this dataset collection can aid clinicians and researchers to study and quickly visualize the functional consequences of a variety of well characterized, naturally occurring mutations on a genome-wide scale.

## Material and Methods

A total of 157 datasets were identified in GEO using the following search query: Homo sapiens AND (“primary immunodeficiency diseases” OR “primary immunodeficiencies diseases” OR PID OR “autosomal recessive” OR “autosomal dominant” OR “inherited deficiency”) AND (“Expression profiling by array”). All GEO entries that were returned with this query were manually curated. This process involved reading all the descriptions available for the datasets, the study designs and corresponding research articles. Finally, a total of 19 datasets were retained because they contained samples which were obtained from well characterized patients with known PIDs (i.e. the genetic etiology had been identified) or the samples were obtained from patients which were considered to have common variable immunodeficiency (CVID). These include datasets that were generated from whole blood, PBMCs, fibroblasts, B cells, iPS and kidney biopsy of individuals with defects in intrinsic and innate immunity, combined immunodeficiencies with associated or syndromic features, autoinflammatory disorders, congenital defects of phagocyte number, functions, or both, predominantly antibody deficiencies and diseases of immune dysregulation. The selected datasets are listed in **Table 1.** A breakdown of the dataset collection by category in accordance to the most recently published update on PID classification from International Union of Immunological Societies Expert Committee [5, 9] is shown in Figure 1.

**Figure 1:**
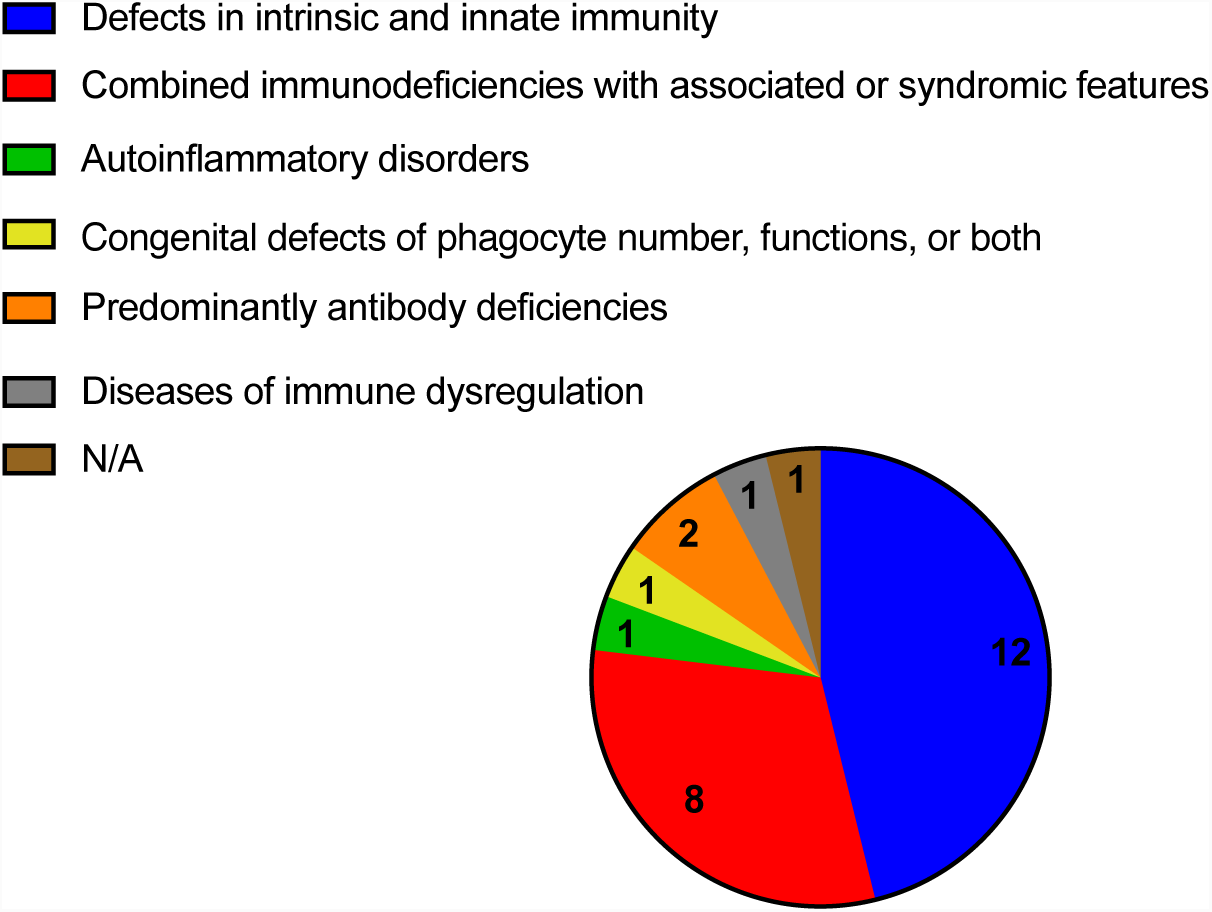
Break down of the dataset collection by category. The pie chart indicates the PIDs classification out for the 19 datasets.

The selected datasets were downloaded from GEO by using the SOFT file format. Then, the datasets were uploaded onto our web tool, called the Gene Expression Browser (GXB), an interactive application hosted on the Amazon Web Services cloud [19]. Information about samples and study design were also uploaded. The available samples were assigned to groups based on the individuals and deficiencies studied and genes were ranked according to different group comparisons allowing the identification of transcripts that were differentially expressed between the patient’s and control subject’s cells cultured or stimulated *ex vivo* under the same conditions. Our dataset collection, uploaded in GXB, is available at http://pid.gxbsidra.org/dm3/geneBrowser/list. A web tutorial for the use of GXB can be accessed at: https://gxb.benaroyaresearch.org/dm3/tutorials.gsp#gxbtut. A detailed description of GXB has been recently published [19-22] and is reproduced here so that readers can use this article as a standalone resource. Briefly, datasets of interest can be quickly identified either by filtering on criteria from pre-defined sections on the left or by entering a query term in the search box at the top of the dataset navigation page. Clicking on one of the studies listed in the dataset navigation page opens a viewer designed to provide interactive browsing and graphic representations of large-scale data in an interpretable format. This interface is designed to present ranked gene lists and display expression results graphically in a context-rich environment. Selecting a gene from the rank ordered list on the left of the data-viewing interface will display its expression values graphically in the screen’s central panel. Directly above the graphical display drop down menus give users the ability: a) To change how the gene list is ranked – this allows the user to change the method used to rank the genes, or to only include genes that are selected for specific biological interest; b) To change sample grouping (Group Set button) – in some datasets, a user can switch between groups based on cell type to groups based on disease type, for example; c) To sort individual samples within a group based on associated categorical or continuous variables (e.g. gender or age); d) To toggle between the bar chart view and a box plot view, with expression values represented as a single point for each sample. Samples are split into the same groups whether displayed as a bar chart or box plot; e) To provide a color legend for the sample groups; f) To select categorical information that is to be overlaid at the bottom of the graph – for example, the user can display gender or smoking status in this manner; g) To provide a color legend for the categorical information overlaid at the bottom of the graph; h) To download the graph as a portable network graphics (png) image. Measurements have no intrinsic utility in absence of contextual information. It is this contextual information that makes the results of a study or experiment interpretable. It is therefore important to capture, integrate and display information that will give users the ability to interpret data and gain new insights from it. We have organized this information under different tabs directly above the graphical display. The tabs can be hidden to make more room for displaying the data plots, or revealed by clicking on the blue “show info panel” button on the top right corner of the display. Information about the gene selected from the list on the left side of the display is available under the “Gene” tab. Information about the study is available under the “Study” tab. Rolling the mouse cursor over a bar chart feature while displaying the “Sample” tab lists any clinical, demographic, or laboratory information available for the selected sample. Finally, the “Downloads” tab allows advanced users to retrieve the original dataset for analysis outside this tool. It also provides all available sample annotation data for use alongside the expression data in third party analysis software. Other functionalities are provided under the “Tools” drop-down menu located in the top right corner of the user interface. Some of the notable functionalities available through this menu include: a) Annotations, which provides access to all the ancillary information about the study, samples and dataset organized across different tabs; b) Cross-project view; which provides the ability for a given gene to browse through all available studies; c) Copy link, which generates a mini-URL encapsulating information about the display settings in use and that can be saved and shared with others (clicking on the envelope icon on the toolbar inserts the URL in an email message via the local email client); d) Chart options; which gives user the option to customize chart labels.

## Data Availability

All datasets included in our curated collection are also available publically via the NCBI GEO website : https://www.ncbi.nlm.nih.gov/gds/ and are referenced throughout the manuscript by their GEO accession numbers (e.g. <u>GSE92466</u>) Signal files and sample description files can also be downloaded from the GXB tool under the “downloads” tab.

## AUTHOR CONTRIBUTIONS

SBouga and NM conceived the theme for this dataset collection. SBouga, NM and SBough, contributed to the query, selection, loading and curation of datasets. DC participated in conceptualization of the approach. SBouga and NM prepared the first draft of the manuscript. All authors were involved in the revision of the draft manuscript and have agreed to the final content.

## ACKNOWLEDGMENTS

We would like to thank all the investigators who decided to make their datasets publically available by depositing them in GEO.

## SUPPLEMENTARY MATERIAL

No supplementary material provided

